# Analysis of splicing abnormalities in the white matter of myotonic dystrophy type 1 brain using RNA-sequence

**DOI:** 10.1101/2022.12.23.521721

**Authors:** Kazuki Yoshizumi, Masamitsu Nishi, Masataka Igeta, Masayuki Nakamori, Tsuyoshi Matsumura, Harutoshi Fujimura, Kenji Jinnai, Takashi Kimura

## Abstract

Myotonic dystrophy type 1 (DM1) is a neuromuscular disorder caused by the genomic expansions of CTG repeats, in which RNA-binding proteins, such as muscleblind-like protein, are sequestered in the nucleus, and abnormal splicing is observed in various genes. Although abnormal splicing reportedly occurs in the brains of patients with DM1, it is relation to the central nervous system symptoms is unknown. Several imaging studies have indicated substantial white matter (WM) defects in patients with DM1. Here, we performed RNA-sequencing and analysis of CTG repeat lengths in the frontal lobe of patients with DM1, separating the grey matter (GM) and WM, to investigate the splicing abnormalities in the DM1 brain, especially in the WM. The results demonstrated the number of repeats in the GM tended to be increase, with several genes showing similar levels of splicing abnormalities in the GM and WM, suggesting that the WM defects in DM1 are not only caused by aberrant splicing of GM RNA but also of WM RNA, which could be attributed to abnormal splicing of glial cell RNAs.

## Introduction

Myotonic dystrophy type 1 (DM1) is the most common type of muscular dystrophy observed in adults. Patients with DM1 have systemic symptoms including skeletal muscle weakness, cardiac conduction defects, cataracts, glucose intolerance, and central nervous system (CNS) dysfunction. Several neurological symptoms, including cognitive impairment, depression, anxiety, and behavioral disorders, have been observed, indicating cerebral involvement [1]. CNS disorders in patients with DM1 result in decreased quality of life [2] and significantly increased disease the burden. DM1 is caused by abnormal expansion of the CTG-trinucleotide repeats in the 3’ untranslated region (UTR) of the *myotonic dystrophy protein kinase* (DMPK) gene [3]. The accumulation of toxic RNAs in the nucleus is caused by mutant CTG repeats in the 3’ UTR of DMPK, which inhibits RNA-binding proteins, causing sequestration of muscleblind-like (MBNL) proteins [4,5] and upregulation of CUGBP/Elav-like family (CELF) proteins [6]. The sequestration of these proteins affects the alternative splicing (AS) of various genes [7]. We previously identified several splicing abnormalities in the brains of MBNL1/2 knockout mice and patients with DM1, demonstrating that the major splicing factor in the brain is MBNL2 and that MBNL1 functions similarly in the skeletal muscles [8,9]. Moreover, combined depletion of MBNL proteins by CUG repeats has been reported as a critical factor in the pathogenesis of DM1 in both the brain and skeletal muscle [10].

In an earlier study, we investigated splicing abnormalities in patients with DM1 based on the brain region and found less mis-splicing in the cerebellum than that observed in other regions [8,11].

Mills et al. employed RNA-sequencing (RNA-seq) to demonstrate that the grey matter (GM) and white matter (WM) of the human frontal lobe show different transcriptome profiles and distinct AS [12]. Neuroimaging analyses of the brains of patients with DM1 showed abnormalities in both the GM and WM [13]. Conventional magnetic resonance imaging has demonstrated abnormalities in both the GM and WM, including reduced GM volume [14], WM hyperintense lesions [15], and anterior temporal white matter lesions (ATWML) [13]. Moreover, voxel-based morphometry (VBM) revealed significant reductions in the GM and WM volumes of patients with DM1 than in the controls [16]. Diffusion tensor imaging studies have revealed a nearly symmetrical reduction of fractional anisotropy in the major association, commissural, and projection fiber tracts, suggesting extensive WM defects [14].

Considering the extensive damage to the GM and WM in the images, we examined whether the splicing regulation of several genes varied between the GM and WM, and found significantly more splicing changes in the GM than in the WM [11].

However, only 15 genes were analyzed in this study. Additionally, transcriptome analysis of the frontal cortex of the DM1 brain, using RNA-seq, revealed new splicing abnormalities in genes such as *GRIP1* [17]. In this case, transcriptome analysis was only performed for the GM; therefore, a comprehensive survey of splicing abnormalities in the GM and WM is necessary.

We aimed to thoroughly investigate the splicing patterns in the GM and WM using RNA-seq to identify WM-specific splicing abnormalities in this study.

## Materials and methods

### Ethics and written informed consent

This study was approved by the Ethics Committee of Hyogo Medical University (Approval No. 93), and written informed consent was obtained from the patients or their families for autopsies.

### Human RNA collection

RNA was extracted from the frontal lobes of autopsied brain samples from five patients with DM1 and five patients with amyotrophic lateral sclerosis (ALS: disease control). The clinical characteristics of each patient are shown in Table 1.

**Table 1.** Clinical characteristics of each patient.

| Disease | Sex | Age (years) | Onset age (years) | Duration (years) | Type of DM1 | Cognitive decline |
| --- | --- | --- | --- | --- | --- | --- |
| DM1 | F | 58 | 38 | 20 | Adult onset | Mild |
|  | M | 47 | 12 | 35 | Juvenile | Severe |
|  | M | 41 | 5 | 36 | Juvenile | Mild |
|  | M | 51 | 40 | 11 | Adult onset | Mild |
|  | M | 57 | 18 | 39 | Juvenile | Mild |
| **ALS** | F | 53 | N/A | N/A | N/A | Normal |
|  | M | 69 | N/A | N/A | N/A | Normal |
|  | F | 73 | N/A | N/A | N/A | Normal |
|  | F | 73 | N/A | N/A | N/A | Normal |
|  | M | 70 | N/A | N/A | N/A | Normal |

Type of DM1; Juvenile onset (age of onset 10 – 20 years), Adult onset (age of onset 20 – 40 years), Late onset (age of onset > 40 years). The degree of cognitive decline was rated on three levels: normal, mild, and severe.

The frontal lobe tissue was sectioned into 60-μm thick sections using a Leica® CM1520 Cryostat (Leica Biosystems, Germany). The boundary between the GM and WM was visually determined, and the tissue was manually separated into GM and WM. RNA was isolated using RNeasy® Plus Mini (QIAGEN^®^, Germany).

To confirm the separation of the GM and WM, we performed quantitative realtime (RT) polymerase chain reaction (PCR) (qPCR), using primers for *NEFH* (expressed to a higher level in the GM), *MOG* (expressed to a higher level in the WM), and *GAPDH* (control) [12] (Table 2), with PowerUp™ SYBR™ using the Thermal Cycler Dice^®^ Real Time System III (TAKARA Bio, Japan). These PCRs were run in triplicates, and the expression levels of *NEFH* and *MOG* were calculated using the 2-ΔΔCt method with GAPDH as a control; the data are shown as the mean of triplicates.

**Table 2.**
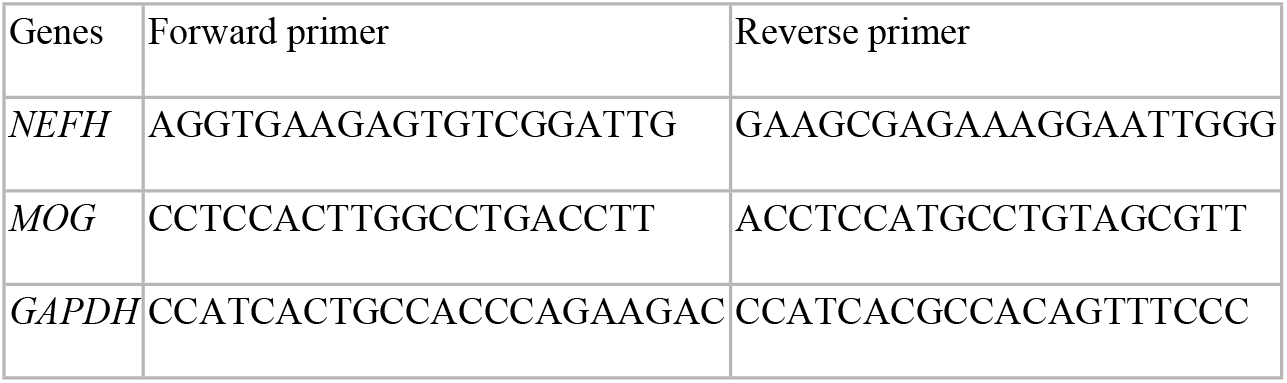
Sequence of primers created for qPCR.

### RNA-seq analysis

The RNA quality was assessed using an Agilent 2100 Bioanalyser (Agilent Technologies, USA). Three samples each of ALS and DM1 were selected for RNA-seq, and their RNA integrity number (RIN) values were >6.0, while the RIN values of the remaining two ALS and DM1 were >3.0. RNA-seq was performed using DNBSEQ-G400 (MGI Tech, Co., Ltd, China) at BGI Genomics and sequenced to a depth of approximately 40 million read pairs. The read quality was checked using fastQC (v.0.11.9). The reads were aligned to the human reference genome (GRCh38) using HISAT2 (v2.0.4) [18]. The aligned and sorted bam files were used for AS analysis using rMATS (v4.1.2) [19]. We calculated the percent spliced in parameter (PSI), denoted as the ratio of alternative exons, using the RNA-seq results.

### Establishing criteria for abnormal splicing

The AS events were identified using the following criteria: a false discovery ratio (FDR) of ≤ 0.05, inclusion and exclusion reads ≥ 20, P value < 0.01 using a likelihood-ratio test, and an average level of PSI (ΔPSI) ≥ 0.2.

### Primer design

Based on the RNA-seq results, the DNA sequences of the splicing events were identified using Genetyx^®^-Network version 16.0.1 (Genetyx Corporation, Japan) and the National Center for Biotechnology Information (NCBI) database. Eight candidate splicing events were abnormal only in the WM and three that were abnormal in both the GM and WM; one of them was *ADD1* exon8, which was examined in our previous study [11]. Ten mis-spliced exons were identified, including *ASPDH* exon 5(150 nt.), *CACNA1A* exon 45(36 nt.), *DNM1* exon 22(37 nt.), *GPM6A* exon 2(59 nt.), *IP6K2* exon 2(139 nt.), *MBD1* exon 9 (75 nt.), *NPRL3* exon 5 (75 nt.), *PALM* exon 8(132 nt.), *PLPP1* exon3(155nt.), and *ZFYVE21* exon6 (54 nt.). Forward and reverse primers were designed for each splicing gene using NCBI Primer-BLAST (https://www.ncbi.nlm.nih.gov/tools/primer-blast/) and Primer3 (https://bioinfo.ut.ee/primer3-0.4.0/) (Table 3).

**Table 3.**
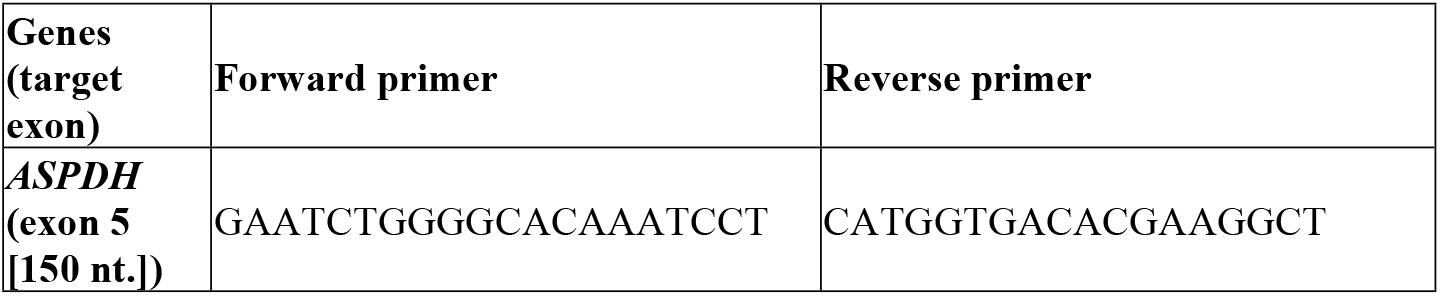
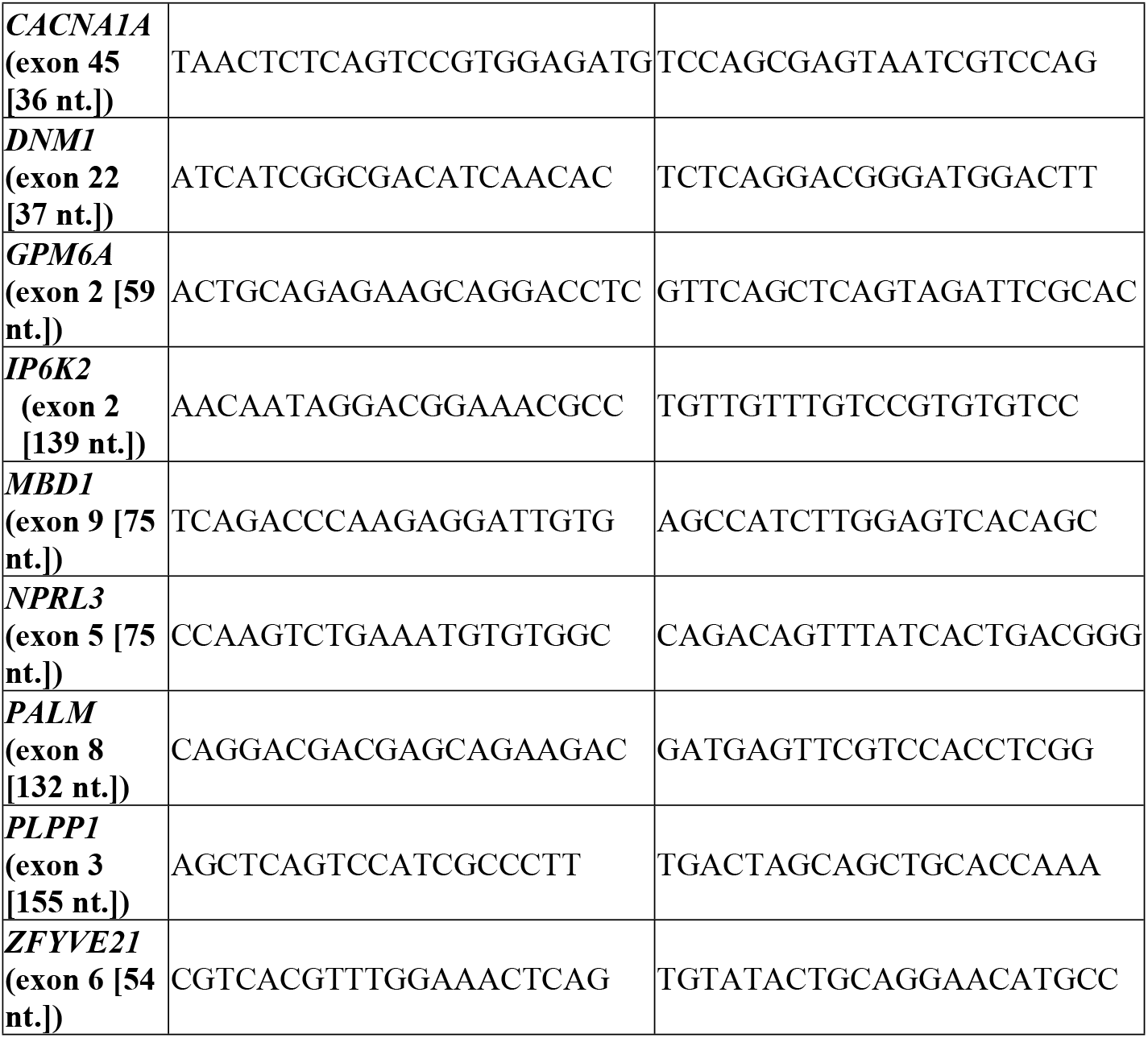
Sequence of primers created for RT-PCR.

### RT-PCR methods

We used random hexamers with the SuperScript® III First-Strand Synthesis System (Invitrogen™, USA) in accordance with the manufacturer’s instructions and synthesized cDNA from 400ng of extracted of RNA. AmpliTaq Gold® 360 Master Mix (Applied Biosystems, USA) was used to amplify cDNA with primers under the following PCR conditions: 94°C for 4 min, 36 cycles of 94°C for 30 s, 61°C for 30 s, and 72°C for 1 min. We analyzed the PCR products using an Agilent 2100 Bioanalyzer.

### Human DNA collection

DNAs, which was also used for RNA-seq, was extracted from the brains of three patients with DM1 and separated into GM and WM using the same method as for RNA. We isolated the DNA by phenol/chloroform extraction and ethanol precipitation as previously described [20].

### Southern blot

Southern blotting was performed as described by Nakamori et al. [21] with slight modifications. Genomic DNA was digested using HaeIII and AluI. The fragments were separated on 0.7% agarose gels buffered with 40 mM Tris-acetate at 4V/cm for 6 h and transferred overnight to nylon membranes (Roche, Switzerland) by alkaline transfer. After UV crosslinking, blots were hybridized with 10 pmol/ml DIG-labeled (CAG)_7_ (5′-gcAgCagcAgca-3′) at 70°C for 4 h in hybridization buffer (5 × SSC, 1% block solution [Roche, Switzerland], 0.1% N-laurylsarcosine, 0.02% sodium dodecyl sulfate). Blots were washed with high stringency (0.5× SSC, 70°C) and developed with alkaline phosphatase-conjugated anti-DIG antibody (Roche, Switzerland) with CDP-Star substrate in accordance with the manufacturer’s instructions, followed by chemiluminescence detection using ChemiDoc MP imager (Biorad, USA).

### Statistical analysis

We performed Welch’s t-test to test the mean difference in the PSI between ALS and DM1, using a two-sided significance level of 5% for all comparisons. We did not adjust the significance level for multiplicity to maintain the maximum power, as indicated by Saville [22]. The average PSI change (ΔPSI) was calculated by subtracting the average PSI of the ALS from DM1.We demonstrated the summary statistics, 95% confidence intervals derived from the unpooled variance, and p-values obtained by Welch’s t-test for the difference between the mean ΔPSI GM and the mean ΔPSI WM for each exon as described in a previous study [11].

To compare the extension of CTG repeats in DM1 GM and WM, p-values were obtained using a one-sample t-test from the number of GM repeats minus the number of WM repeats in the same patient. This statistical analysis was performed using EZR (Saitama Medical Center, Jichi Medical University, Japan) [23], which is a graphical user interface for R (The R Foundation for Statistical Computing, Austria). The number of peak repeats was considered as the number of CTG repeats.

## Results

### Confirmation of separation of the GM and WM

qPCR analysis revealed that the expression levels of *NEFH* to be higher in the GM samples, whereas those of *MOG* were higher in the WM, verifying that each sample was successfully segregated into GM and WM (Fig 1).

**Fig 1.**
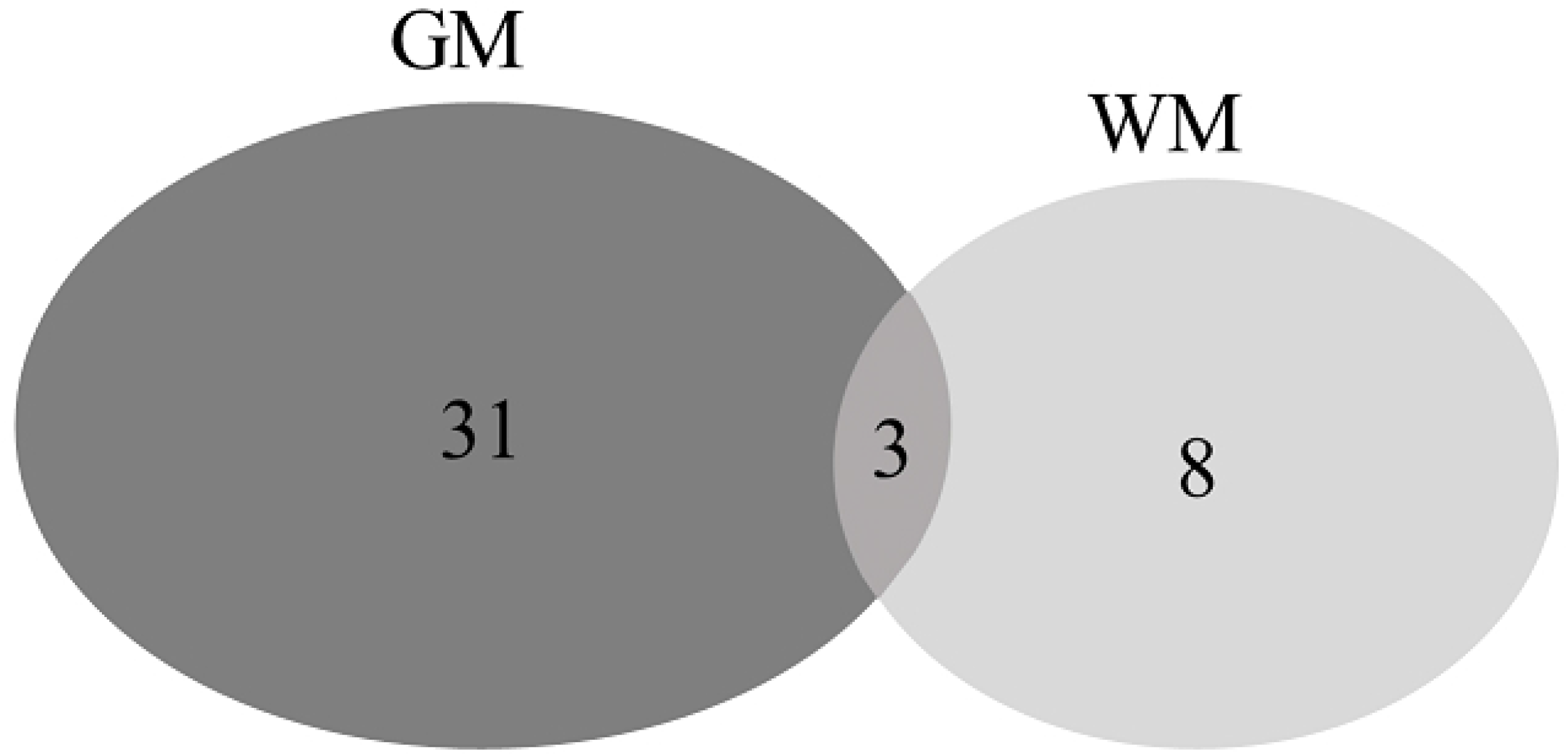
Comparison of the *NEFH* and *MOG* expression levels in the GM and WM. The expression levels of *NEFH* and *MOG* were calculated using the 2-ΔΔCt method with *GAPDH* as a control, and data are shown as the mean of triplicates. GM: Grey matter, WM: White matter.

### Comprehensive study of mis-splicing with RNA-seq

We identified 31 mis-spliced events in the GM, eight events in the WM, and three events in both GM and WM using our criteria (Fig 2). These WM events included *PALM* exon 8 and *ZFYVE21* exon 6, which were previously reported to be mis-spliced in the frontal cortex and heart of DM1, respectively [17,24].

**Fig 2.**
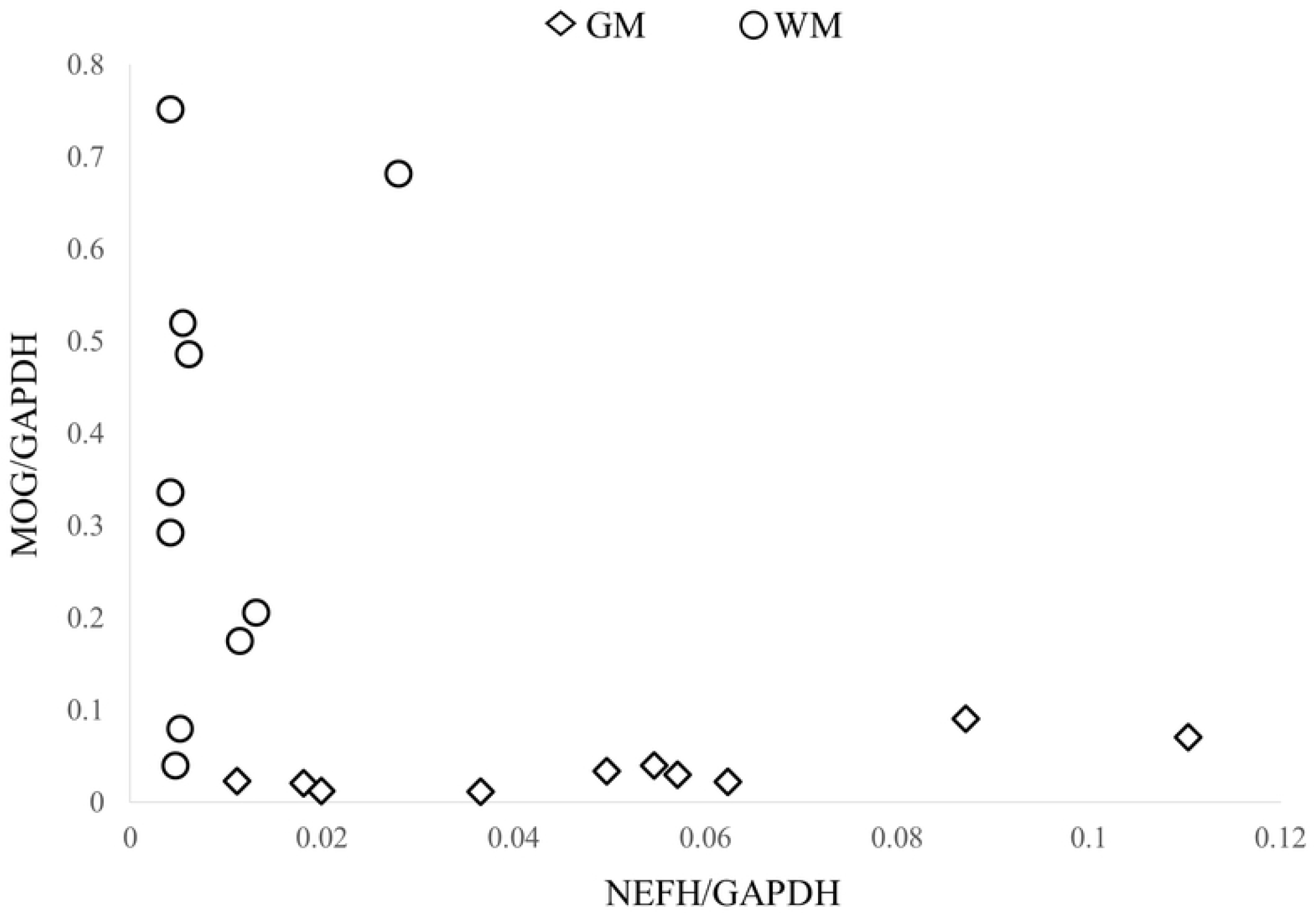
Number of mis-spliced genes in the GM and WM found by RNA-seq. This Venn diagram shows the number of abnormally spliced exons in the GM and WM of myotonic dystrophy type 1 (DM1) under the defined criteria. The overlapping areas in the Venn diagram are genes that showed abnormal splicing in both the GM and WM. GM: Grey matter, WM: White matter.

### Validation of mis-splicing with RT-PCR

To validate the splicing abnormalities in the WM, ten splicing events in the WM (excluding *ADD1* exon 8) were examined by RT-PCR in five ALS and five DM1 brains, including three ALS and three DM1 brain samples that were used in the RNA-seq study.

Compared to the ALS brains, the PSI of *PALM* exon 8, *CACNA1A* exon 45, *ZFYVE21* exon 6, and *MBD1* exon 6 in the GM and WM of the DM1 brains were significantly different, suggesting that these genes were spliced in both the GM and WM of DM1 brains (Fig 3). The PSI of *PLPP1* exon 2 was significantly different in the GM, but not in the WM of DM1 brains compared to ALS brains. The PSI of *DNM1* exon 22 could not be calculated owing to the nonspecific PCR products (data not shown).

**Fig 3.**
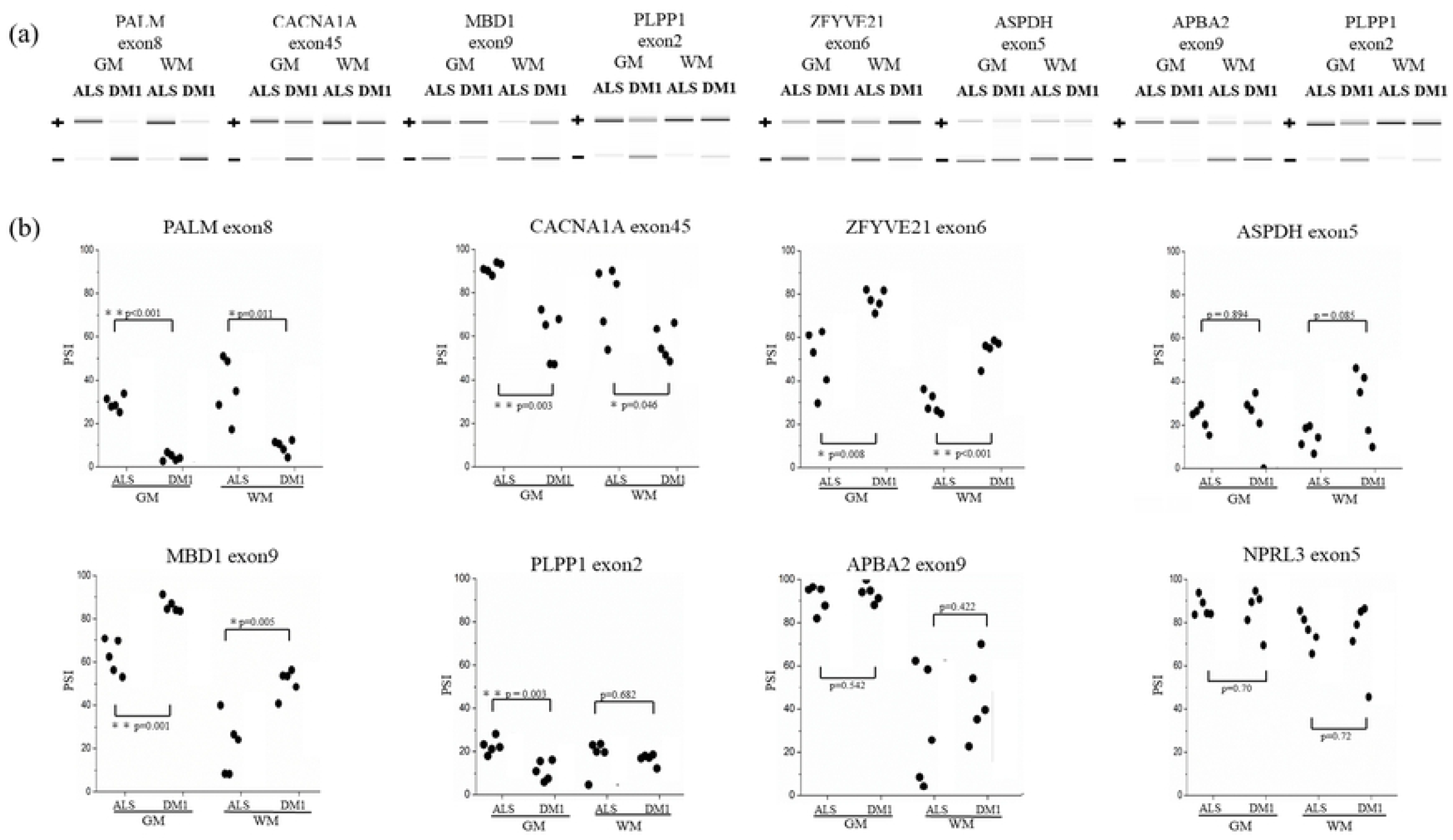
Validation of mis-splicing by RT-PCR. (a) RT-PCR products from the GM and WM in ALS and DM1. (b) PSI of alternative exons in the GM and WM for candidate genes from RNA-seq results. The PSI values for each gene were analyzed using Welch’s t-test. The graph is shown as a box and-whisker diagram, with the line in the box representing the median, and the square symbol representing the mean. RT-PCR: real-time polymerase chain reaction, PSI: Percent spliced in parameter, ALS: Amyotrophic lateral sclerosis, DM1: Myotonic dystrophy type 1, GM: Grey matter, WM: White matter.

Next, ΔPSI was used to compare the extent of mis-splicing of the exons that showed a significant difference between the DM1 and ALS brains, between the GM and WM (Fig 4). The mean PSI of *PALM* exon 8 was 29.26% and 36.12% for the GM and WM in ALS brains, respectively, and 4.36% and 9.23% for the GM and WM in DM1 brains, respectively. The ΔPSI of the GM was –24.90%, and that of the WM was –26.89% between the two disorders. The difference in the ΔPSI between DM and ALS [95% confidence intervals] was –1.98%, [–19.282%, 15.318%], and the *p*-value was 0.7735. Among all the five examined samples, no significant differences in ΔPSI were found between the GM and WM, suggesting that abnormal splicing of these genes in the GM and WM occurs to the same extent.

**Fig 4.**
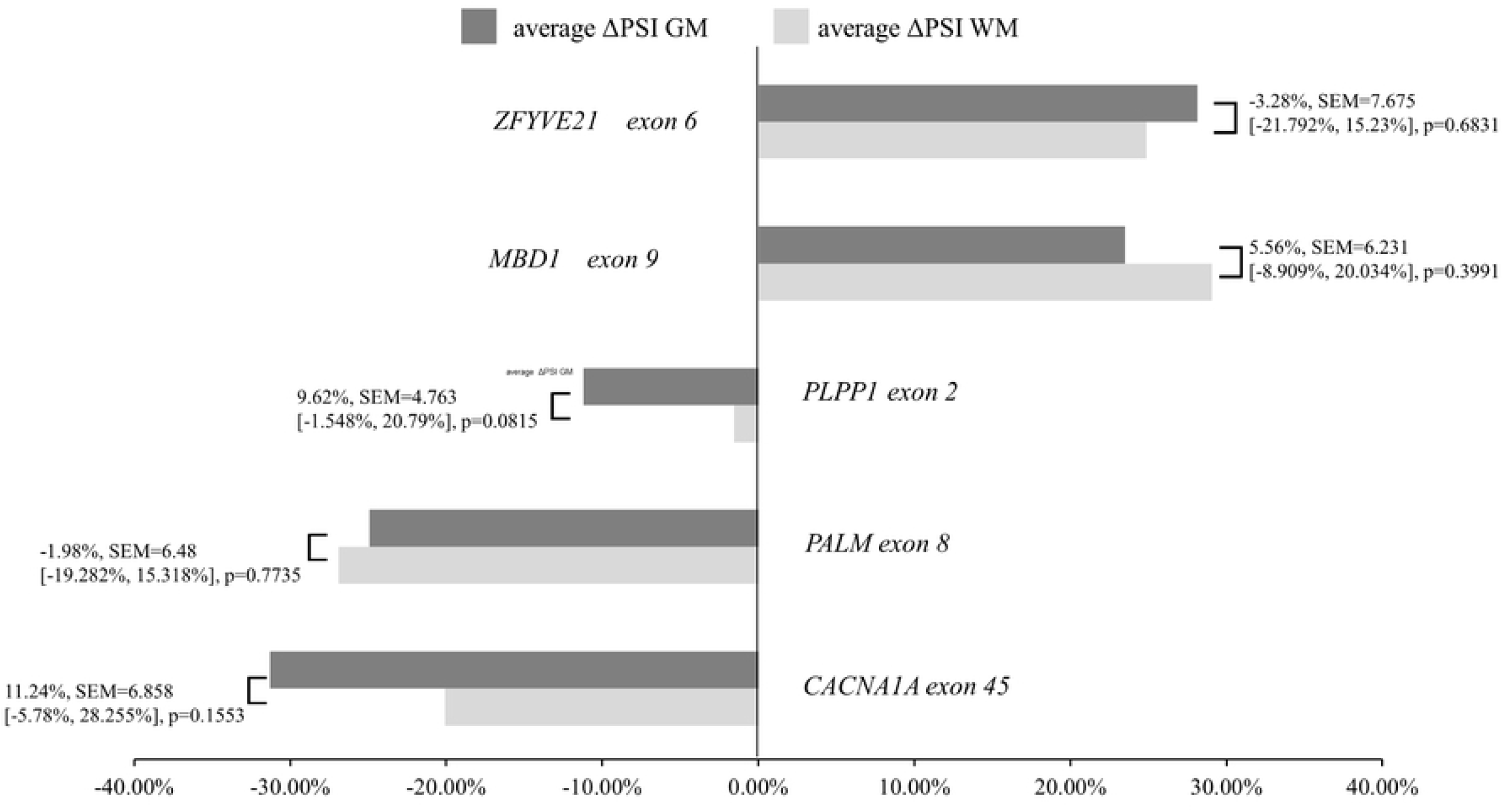
Comparison of the degree of mis-splicing in the GM and WM using ΔPSI. Bar graph showing a comparison of the mean ΔPSI GM and mean ΔPSI WM of five samples using Welch’s t-test for genes that showed mis-splicing in the GM and WM of in DM1 brains by RT-PCR. The numbers in the graphs represent the mean difference for comparison of the mean ΔPSI GM and mean ΔPSI WM, SEM, [Lower CL, Upper CL], and p-values in this order. RT-PCR: real-time polymerase chain reaction, PSI: Percent spliced in parameter, DM1: Myotonic dystrophy type 1, GM: Grey matter, WM: White matter.

### Comparison of the CTG repeat lengths in the GM and the WM of DM1

The peak density of the longest band for each sample was estimated based on the expansion size. The number of CTG repeats in each sample was as follows (GM, WM): (1) 2970, 2370; (2) 3060, 2890; and (3) 3150, 2510 (Fig. 5), with no statistically significant difference between the GM and WM (P = 0.089, one-sample t-test); however, the number of repeats in the GM tended to be extended in all the samples.

**Fig 5.**
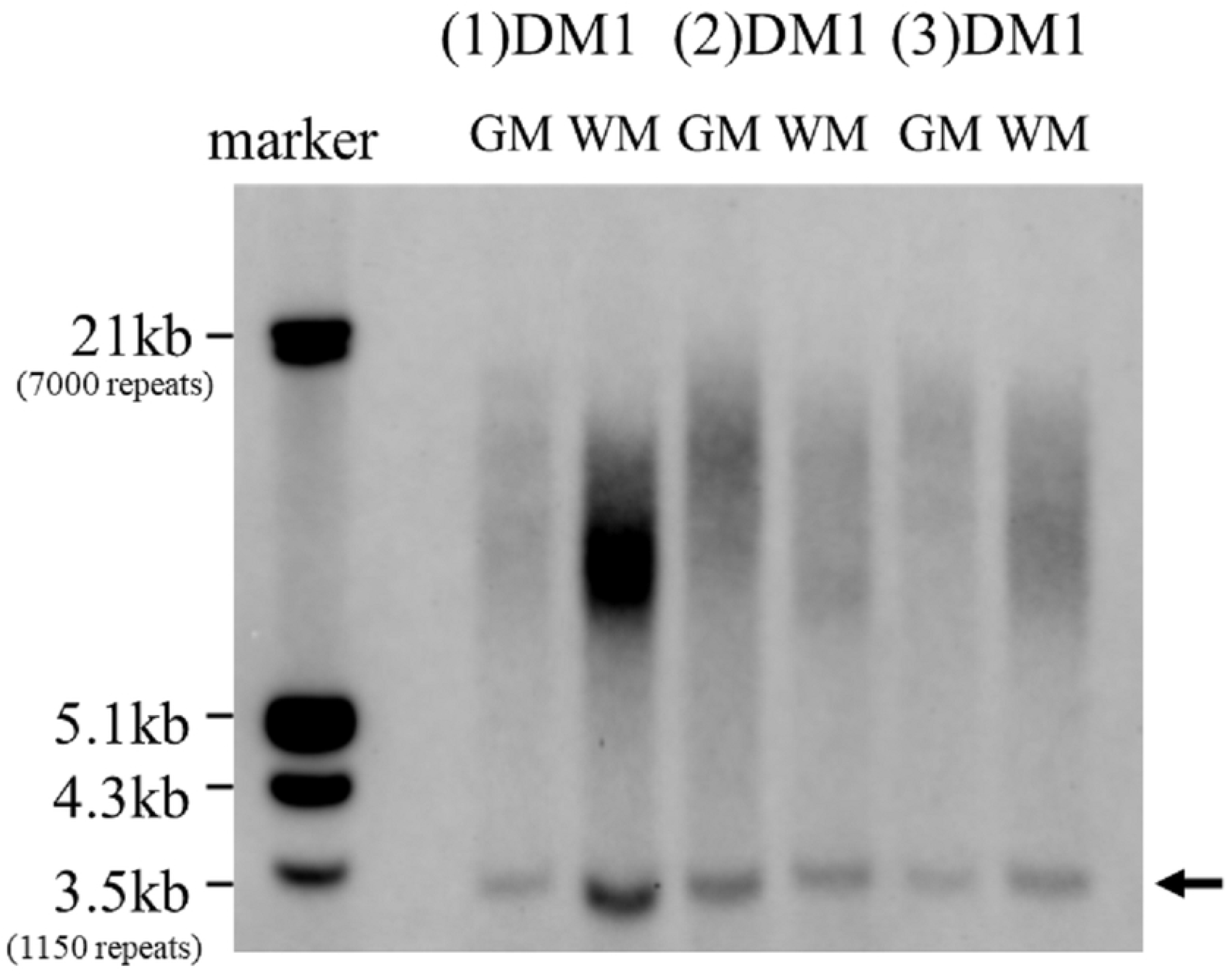
Southern blot analysis of DNA from the GM and WM in DM1 brains. From the left lane, molecular weight markers, (1) DM1: GM WM, (2) DM1: GM WM, and (3) DM1: GM WM. The arrow indicates nonspecific signal. The peak density of the longest band for each sample was estimated based on the expansion size. The number of CTG repeats was as follows (GM, WM): (1) 2970, 2370; (2) 3060, 2890; (3) 3150, 2510. DM1: Myotonic dystrophy type 1, GM: Grey matter, WM: White matter.

## Discussion

In the present study, RNA-seq was performed on the GM and WM tissue samples of the brains of patients with DM1. A greater number of mis-spliced genes were found in the GM; however, mis-splicing was also observed in several genes in the WM. Although RT-PCR validation did not reveal any genes causing abnormal splicing in the WM only, the following genes in the WM showed the same level of mis-splicing as in the GM: *PALM* exon 8, *CACNA1A* exon 45, *ZFYVE21* exon 6, and *MBD1* exon 6. Of these four genes, three (*MBD1* exon 9, *CACNA1A* exon 45, and *PALM* exon 8) have been reported as being in the top 130 high-confidence splicing events in the frontal cortex of the DM1 brain by Otero et al. [17]. These three previously reported genes, plus one newly identified gene, *ZFYVE21* exon6, were mis-spliced to the same extent in the WM. Therefore, we suggest that these splicing changes might be related to substantial WM damages in DM1 brains.

We previously examined the differences in the splicing regulation of several genes between the GM and WM and reported significantly more splicing abnormalities in the GM than in the WM. In seven of the 15 genes examined, the extent of splicing change between DM1 and ALS was significantly increased in the GM compared to that in the WM. In contract, none of the genes showed a significant increase in the extent of splicing change in the WM versus the GM.

The results of RNA-seq and RT-PCR revealed four new genes, *PALM* exon 8, *MBD1* exon 9, *CACNA1A* exon 45 and *ZFYVE21* exon 6, which showed significant abnormal splicing in the WM.

*PALM* localizes to axons, the plasma membranes of postsynaptic specializations, and dendritic processes. Alternative splicing of *PALM* exon 8 modulates the reduction in synapse formation, filopodia induction and AMPA receptor recruitment [25]. These changes are associated with alterations in synaptic and structural plasticity, including the induction of long-term potentiation. *MBD1* is involved in epigenetic gene regulation via DNA methylation [26]. Since Mbd1–/– mice exhibit symptoms of reduced social interaction, learning disabilities, anxiety, sensorimotor gating deficits, and depression, *MBD1* could be associated with these symptoms [27]. Some of these symptoms are similar to the CNS symptoms observed in patients with DM1. However, the effects of alternative splicing of *MBD1* exon 9 have not yet been reported.

No previous study has reported on the effects of alternative splicing of *CACNA1A* exon 45 and *ZFYVE21* exon 6, both of which were shown to be mis-spliced in the GM and WM in this study. The *CACNA1A* gene encodes the P/Q-type voltagegated Ca2+ channel α1 subunit CaV2.1 and is involved in neurotransmitter release and synaptic plasticity [28,29]. *CACNA1A* has been reported to be associated with episodic ataxia type 2, familial hemiplegic migraine type 1, and spinocerebellar ataxia type 6 [30]. *ZFYVE21* has been reported to be a complement inducer of NF-κB in endothelial cells [31]; however, its function in the nervous system is poorly understood. Mis-splicing of these genes could possibly result in changes in the WM and leading to CNS symptoms.

We have added new insights to our previous discussion of splicing differences between the GM and WM [11] and proposed several explanations for our findings. First, the cell types vary between the GM and WM; neuronal cell bodies, protoplasmic astrocytes, and microglial cells are found primarily in the GM, while the WM includes axons, oligodendrocytes, fibrous astrocytes, microglial cells, and ependymal cells. In the DM1 brains, although the effects on non-neuronal cells remain unclear, RNA foci accumulate in the astrocytes, oligodendrocytes, and neurons [32,33]. Dincã et al. reported that astrocytes in the frontal cortex and hippocampus of mice are affected by CUG-RNA toxicity, and mis-splicing, which is observed in the astrocytes but not in the neurons, could be related to the functional defects in cell adhesion and spreading [34]. Accumulation of RNA foci and splicing abnormalities have also been reported in the oligodendrocytes of mice [35]. Many mis-splicing events have been found in the WM glial cells of patients with DM1, and some of the mis-splicing events are more prominent in the WM glial cells than in the GM neurons [36]. Thus, splicing abnormalities in the glial cells, such as astrocytes and oligodendrocytes, could be related to the differences in splicing defects between the GM and WM. Second, it is possible that different transcriptional activity in the axons and neuronal cell bodies could possibly affect the distribution of splicing isoforms. Some mRNAs are transcribed by RNA-binding proteins in the neuronal cell body and then transported to the axons [37]. Protein synthesis in the axons, termed local protein synthesis, occurs primarily in developing axons and during regeneration of injured axons [38]. The distribution of mRNAs in axons was examined using transcriptome analysis [39]. Axons in early development have a propensity to express mRNAs related to axon extension, while those in late development tend to express mRNAs encoding proteins related to dendrites and synapse formation. The selectivity of transported mRNAs could be linked to the splicing differences between the GM and WM. Third, the difference in the number of repeats between the GM and WM could propel abnormal splicing. Indeed, Nakamori et al. measured the CTG repeat length in the neurons of the GM and glial cells of the WM in the anterior temporal lobe and reported that neurons of the GM had significantly longer repeats [36]. In the present study, the number of repeats in the GM and WM was not statistically significantly different; however, the number of repeats in the GM was extended in all the samples. This difference in the number of repeats could explain the differences in splicing.

It remains unclear how differences in splicing between the GM and WM are linked to brain damage. In WM pathology, loss of myelin, widened perivascular spaces in deep and subcortical WM, and loss of axons have been reported; however, some reports indicated that axons remain relatively intact [40]. *PALM* exon8, which was found to have splicing defects in this study, has been previously reported to be mis-spliced in mouse astrocytes [35], suggesting that differences in the splicing isoforms in glial cells affect the splicing defects between the GM and WM. We had predicted Wallerian degeneration as the cause of this WM damage owing to more abundant splicing abnormalities in the GM. However, based on the results of the current study and the preservation of axons in some cases in pathology, we considered the possibility that the WM disorder of the brain in DM1 to be affected by splicing abnormalities in the glial cells, including astrocytes and oligodendrocytes. The genes that were newly identified to be aberrantly spliced in the WM in this study could be related to the WM disorders in the brain of patients with DM1.

## Limitations

This study has several limitations. First, we used ALS brain samples for disease control, which could show splicing changes in the GM and WM [41]. However, we previously compared the splicing between disease controls (7 of 9 samples were ALS) and healthy controls and found no significant difference [9]. Second, the sample size used for RNA-seq was limited. Since Otero et al. included 21 patients with DM1 in their RNA-seq analysis [17], we need to validate these findings using more samples in the future. Finally, our WM samples contained not only axons but also various cells, such as glia cells; therefore, it may be necessary to isolate individual cells using laser microdissection and perform RNA-seq to account for the heterogeneity of the sample.

## Conclusion

In this study, we performed RNA-seq and found several genes that revealed splicing defects in the WM and GM of DM1 brains. In particular, abnormal splicing of *PALM* exon 8 has been previously reported in mouse astrocytes. These results suggest that splicing abnormalities in the astrocytes and other glial cells, may be involved in WM defects in the DM1 brain.

## Acknowledgements

This work was supported by JSPS KAKENHI Grant Number JP18K07515.

We thank editage (www.editage.jp) for English language editing.

## Conflicts of interest

There are no conflicts of interest.

## Notes

### Competing Interest Statement

The authors have declared no competing interest.

